# *In vivo* visualization of intracellular pH gradient in brain using PET imaging

**DOI:** 10.1101/2023.06.21.546029

**Authors:** Tomoteru Yamasaki, Wakana Mori, Takayuki Ohkubo, Atsuto Hiraishi, Yiding Zhang, Yusuke Kurihara, Nobuki Nengaki, Hideaki Tashima, Masayuki Fujinaga, Ming-Rong Zhang

**Affiliations:** Department of Advanced Nuclear Medicine Sciences, Institute for Quantum Medical Science, Quantum Life and Medical Science Directorate, National Institutes for Quantum Science and Technology, 4-9-1 Anagawa, Inage-ku, Chiba, 263-8555 Japan; SHI Accelerator Service Co. Ltd, 7-1-1 Nishigotanda, Shinagawa-ku, Tokyo, 141-0031 Japan

**Keywords:** hydrolysis, hypoxia, intracellular pH, monoacylglycerol lipase, PET

## Abstract

Intracellular pH (pHi) is a valuable index for predicting hypoxic brain damage. However, no positron emission tomography (PET) probe is currently available for monitoring pHi *in vivo*. In this study, we developed a new approach for visualizing monoacylglycerol lipase (MAGL) activity in the brain. This approach used PET with a new probe [^11^C]HC-A, an azetidine carbamate inhibitor, whose uptake and residence depended on the pHi gradient was evaluated with *in silico*, *in vitro*, and *in vivo* assessments. Molecular dynamics simulations predicted that complex (complex-A) between HC-A and MAGL would be difficult to hydrolyze under acidic conditions. *In vitro* assessment using rat brain homogenate showed that [^11^C]HC-A reacted with MAGL to yield [^11^C]complex-A, which was rapidly hydrolyzed to liberate ^11^CO_2_. The ^11^CO_2_ liberation rate was slower at lower pH. In PET with [^11^C]HC-A using ischemic rats, the radioactivity clearance rate, which reflects the production rate of ^11^CO_2_ in the brain, was lower in a remarkably hypoxic area than in the contralateral region. In conclusion, we successfully visualized the pHi gradient in the brain using PET imaging.

## Introduction

Stroke is a major disease involving more than 15 million patients each year worldwide, and its rate is dramatically increasing because of population aging. Cerebral vascular hypoxia damages the central nervous system (CNS) (Freeman & Barone, 2005; Zhang *et al*, 2010; Bădescu *et al*, 2016; Baillieul *et al*, 2017) by switching aerobic energy metabolism to anaerobic metabolism, resulting in intracellular hyperaccumulation of acidic residues such as protons, lactate, and carbonic acid (Bouzat, 2014). Subsequently, the neuronal synapse is depolarized by the disproportion of cations (Na^+^ and K^+^) because the production of adenosine triphosphate (ATP) decreases in anaerobic metabolism. The intracellular Ca^2+^ concentration then increases, and excess neurotransmitters (e.g., glutamate) are released, inducing neurotoxicity, neuroinflammation, and neuronal death (Uria-Avellanal & Robertson, 2014). Such acute damage can lead to a number of pathologies such as depression, post-stroke dementia, and potentially, neurodegeneration (Bădescu *et al*, 2016). Therefore, monitoring changes in intracellular pH (pHi), which depend on the intracellular accumulation of acidic sources, should be very important for predicting neuronal damage.

Positron emission tomography (PET) is an advanced imaging technique widely used in basic and clinical studies to elucidate the kinetics, molecular density, and distribution of drugs *in vivo*. To date, only two major radioprobes have been developed for pH-weighted PET imaging. Four decades ago, [^11^C]dimethyloxazolidinedione ([^11^C]DMO), the membrane permeability of which depends on the pH gradient, was first developed as a pH-weighted marker for PET studies (Kearfott *et al*, 1983). However, PET imaging with [^11^C]DMO was inaccurate because the biodistribution of [^11^C]DMO could not distinguish between plasmalemmal, intracellular, and extracellular pH gradients (Zhang *et al*, 2010). Two decades later, [^64^Cu]DOTA-pHLIP, which specifically binds to the membranes of low-pH cells, was identified (Vāvere *et al*, 2009). However, the binding of [^64^Cu]DOTA-pHLIP depends on the pH gradient, but not on the pHi. To the best of our knowledge, no pH-sensitive PET probe is currently available for use in humans.

Recently, Butler et al. reported azetidine and piperidine carbamates as covalent inhibitors of monoacylglycerol lipase (MAGL), an enzyme widely distributed throughout the mammalian brain, and a key enzyme for regulation of the endocannabinoid system (Butler *et al*, 2017). Our group recently developed radiolabeled azetidine carbamate inhibitors for use as PET probes for imaging of MAGL (Fig 1A). Although they were designed as irreversible PET probes, they showed unexpected rapid radioactivity clearance, similar to that shown by reversible probes, after a high initial uptake of radioactivity in the brain (Cheng *et al*, 2018; Chen *et al*, 2019; Mori *et al*, 2019). Such brain kinetics strongly suggest that carbamate inhibitors covalently bind to MAGL, resulting in unstable complexes in the brain. In this study, we hypothesized a disassociation mechanism for [^11^C]azetidine carbamates in the brain (Fig 1B). [^11^C]Azetidine carbamate covalently reacts with MAGL to form an [^11^C]complex, which is subsequently hydrolyzed, resulting in ^11^CO_2_ as a final radioactive product. Because ^11^CO_2_ is rapidly cleared from the brain (Brooks *et al*, 1984), we considered that the radioactivity clearance rate would depend mainly on the rate of hydrolysis of the [^11^C]complex in the brain.

**Figure 1.**
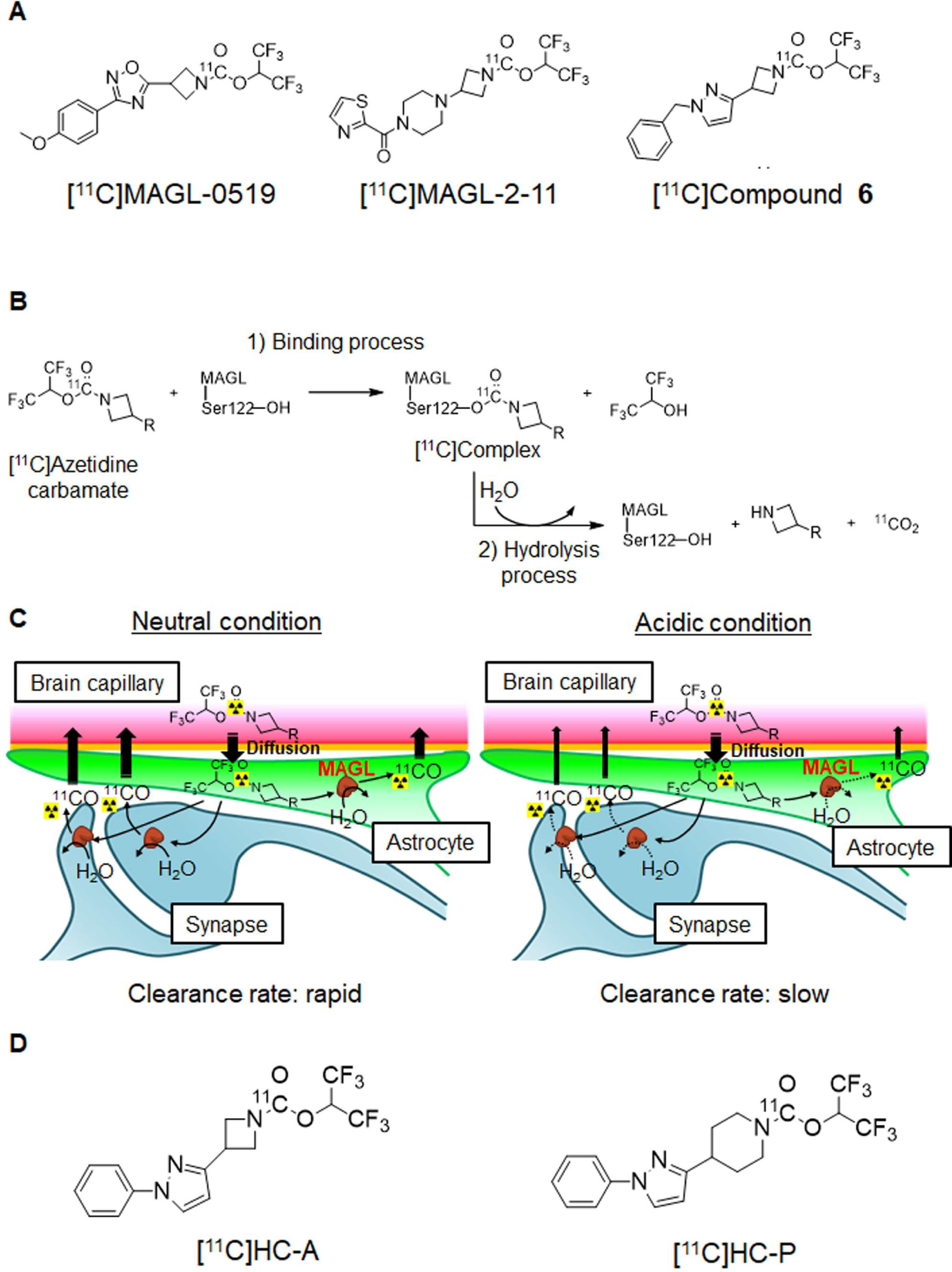
Hypothesis of hydrolysis mechanism for PET probes of covalent azetidine inhibitors for MAGL. A Chemical structures of previous PET probes containing azetidine carbamate ([^11^C]MAGL-0519, [^11^C]MAGL-2-11, and [^11^C]compound **6**) for imaging of MAGL. B, C Proposed disassociation route (B) of the ^11^C-labeled probe containing azetidine carbamate moiety, and a diagram (C) illustrating the effect of acidic sources on ^11^CO_2_ production. D Chemical structures of [^11^C]HC-A and [^11^C]HC-P, novel radioprobes for MAGL imaging developed in this study.

Enzyme activity is reduced at pH values outside the optimal range, generally because changes in pH affect enzyme-substrate complex formation, substrate ionization, and the three-dimensional (3D) structure of enzymes. MAGL is widely expressed throughout the brain and is intracellularly localized not only in pre- and post-synaptic neurons but also in astrocytes (Di Marzo et al, 2015; Viader *et al*, 2015). Therefore, estimation of the ^11^CO_2_ production rate through hydrolysis of the [^11^C]complex formed between azetidine and MAGL may permit monitoring of regional changes in pHi within the brain (Fig 1C).

In this study, we developed 1,1,1,3,3,3-hexafluoropropan-2-yl-3-(1-phenyl-1H-pyrazol-3-yl)azetidine-1-[^11^C]carboxylate ([^11^C]HC-A, IC_50_ = 0.2 nM) as a novel radioprobe for visualizing the pHi gradient (Fig 1D). Simultaneously, we synthesized a new piperidine analog ([^11^C]HC-P, IC_50_ = 2.0 nM) for comparison with [^11^C]HC-A (Fig 1D). We first performed molecular dynamics (MD) simulations for the complexes (defined as complex-A or complex-P) between HC-A or HC-P and MAGL, and then confirmed their structural relationship with *in vivo* hydrolytic instability. Subsequently, we radiosynthesized [^11^C]HC-A and [^11^C]HC-P, and tested their ability to measure the pHi-gradient *in vitro*. To our knowledge, this is the first study to have successfully demonstrated *in vivo* mapping of the pHi gradient based on the production rate of ^11^CO_2_ in ischemic areas of rat brain using PET with [^11^C]HC-A.

## Results and Discussion

### Molecular dynamics simulations

To understand the *in vivo* hydrolytic instability of complex-A and complex-P, we performed MD simulations under neutral conditions similar to normal physiological conditions and compared the results between the two compounds (Figs 2A, B). The root mean square deviation (RMSD) values for complex-A and complex-P showed no significant changes in MD simulations (5 ns) under constant temperature (300 K). Because water is essential for the enzyme hydrolysis of complex-A and complex-P, we further analyzed the accessibility of water molecules to the complexes. By forming hydrogen bonds, His269 and His121 in MAGL were able to capture the water surrounding the carbonyl moiety in both complexes. However, the water molecule was closer to the carbonyl moiety of complex-A (3.4 Å) than to that of complex-P (4.2 Å). Moreover, a hydrogen bond was formed between complex-A and Cys242 in MAGL, which was absent in the case of complex-P because of the longer distance (5.0 Å). These differences indicated that complex-A was hydrolyzed more easily than complex-P. Moreover, the structural strain of the azetidine skeleton is larger than that of the piperidine skeleton, leading to easier hydrolysis.

**Figure 2.**
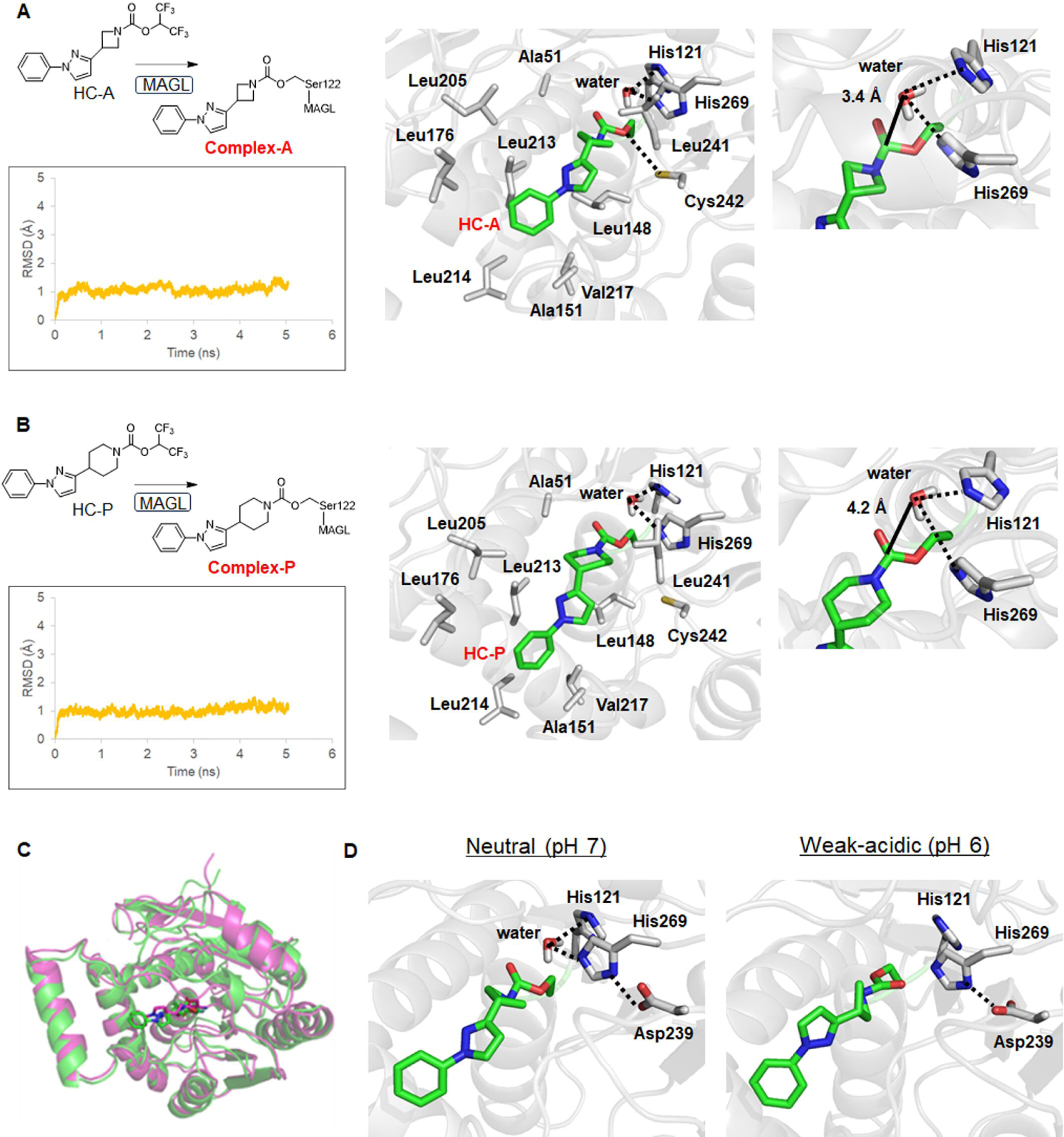
Molecular dynamics (MD) simulations. A, B Chemical structures of HC-A, HC-P, and their MAGL complexes, charts of RMSD values for main-chain atoms of the complexes (A: complex-A; B: complex-P) in MD simulations (5 ns) under neutral conditions, and three-dimensional (3D) structures of the complexes. Hydrogen bonds in the complexes are shown as dotted lines. MD simulations were performed using 3D structures of MAGL (PDB: 6AX1). C Overlaid structures of complex-A composed from MD simulations at pH 7 and pH 6 (green: pH 7; magenta: pH 6). D Changes in the interaction of complex-A with water molecules between neutral (pH 7) and weak-acidic (pH 6) conditions. Spatial localization and interaction of the residue of complex-A with a water molecule at pH 7 (left). Changes in the interaction between the residue of complex-A and a water molecule at pH 6 (right).

As mentioned above, enzyme activity is generally weakened by divergence from the optimum pH. In the present study, we also compared the interaction of complex-A with water molecules at pH values between neutral (pH 7) and weakly acidic (pH 6). The RMSD value calculated by overlaying the final structures of the MAGL main chain between pH 6 and pH 7 was 1.9 Å (Fig 2C), suggesting that the *in vivo* binding affinity of HC-A for MAGL would not change at least at pH 6. By estimating the distance between the carbonyl moiety and the water molecule, we found that the imidazole ring in His269 of MAGL could form a hydrogen bond with water molecules under neutral conditions (Figure 2D). However, at pH 6, water molecules would be difficult to access the carbonyl moiety because of being not captured by the imidazole ring in MAGL, thus enhancing the stability of complex-A against hydrolysis. Moreover, His269 in MAGL maintained a strong electronic interaction with Asp239 in MAGL under weakly acidic conditions. This was possibly due to the protonation of the imidazole ring in His269 under such conditions, which would also hinder the hydrolysis of complex-A.

Taken together, the MD simulations predicted that HC-A covalently bound to MAGL, and that the resulting complex-A was easily hydrolyzed *in vivo* because of the close spatial arrangement between the carbonyl group of complex-A and water molecules. Furthermore, the hydrolysis rate of complex-A would be delayed under acidic conditions because the accessibility of the carbonyl moiety in complex-A to water molecules would be hindered.

### Radiochemistry

We therefore performed *in vitro* and *in vivo* studies to validate our proposed pHi gradient detection approach shown in Fig 1C. We radiosynthesized [^11^C]HC-A and [^11^C]HC-P based on our previous report, using ^11^COCl_2_ as a labeling agent (Wang *et al*, 2016). Briefly, ^11^COCl_2_ gas was trapped in a solution containing 1,1,1,3,3,3-hexafluoro-2-propanol and 1,2,2,6,6-pentamethylpiperidine (PMP) to form [^11^C]chloroformate as an intermediate, which was reacted with each precursor to produce [^11^C]HC-A or [^11^C]HC-P (Fig 3A). Starting with 30 GBq of ^11^CO_2_, [^11^C]HC-A and [^11^C]HC-P were synthesized with 0.9 ± 0.5 GBq (3.1 ± 1.5%) and 1.1 ± 0.6 GBq (4.3 ± 2.3%) at the end of synthesis (EOS), respectively. The radiochemical purities of [^11^C]HC-A and [^11^C]HC-P were > 98%, and the molar activities were 101 ± 24 GBq/μmol for [^11^C]HC-A (n = 11) and 62 ± 13 GBq/μmol for [^11^C]HC-P (n = 5) at the EOS. No radiolysis was observed up to 90 min after formulation, suggesting the radiochemical stability of [^11^C]HC-A and [^11^C]HC-P over the duration of at least one PET scan.

**Figure 3.**
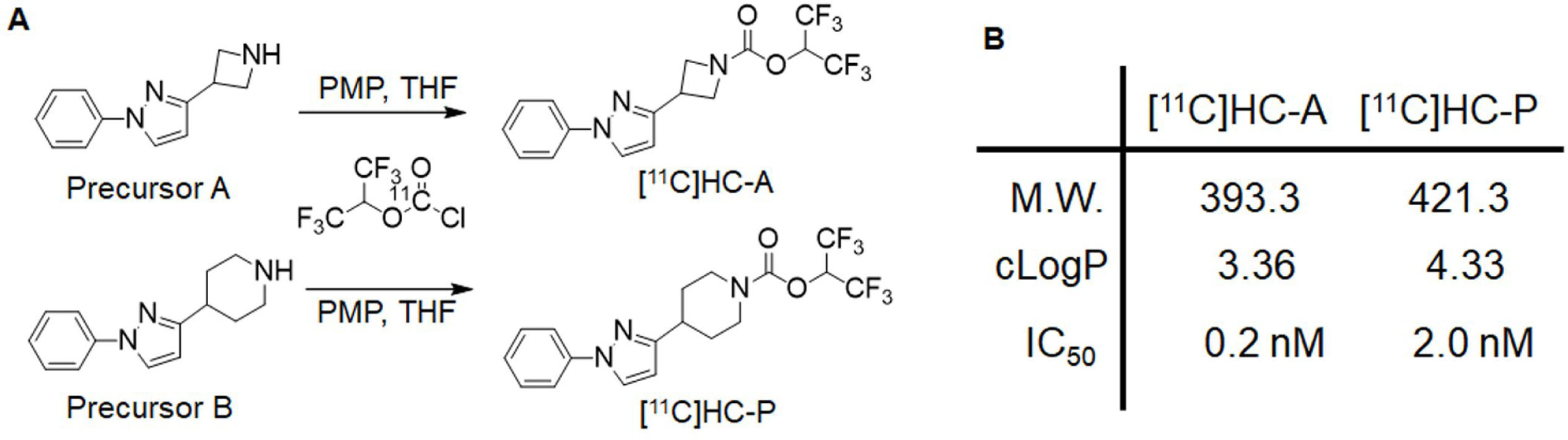
Radiosynthesis of [^11^C]HC-A and [^11^C]HC-P. A [^11^C]HC-A and [^11^C]HC-P were synthesized by reacting respective precursors (A for [^11^C]HC-A, B for [^11^C]HC-P) and [^11^C]chloroformate in a solution containing 1,2,2,6,6-pentamethylpiperidine (PMP) and THF. B General profiles (molecular weight, lipophilicity, and a half maximal inhibitory concentration for MAGL) of [^11^C]HC-A and [^11^C]HC-P.

### *In vitro* ^11^CO_2_ collection assay

Next, to confirm the production of ^11^CO_2_ resulting from MAGL hydrolysis, we conducted an *in vitro* ^11^CO_2_ collection assessment (Barzagli *et al*, 2016) of [^11^C]HC-A or [^11^C]HC-P using rat brain homogenate (Fig 4A). Fig 4B shows the radioactivity expressed as a percentage of the injected dose (%ID) in three divided compartments (C1: unbound radioprobe; C2: complex with MAGL; C3:^11^CO_2_ as a final product). In the C1 compartment representing the unreacted radioprobe, the radioactivity of [^11^C]HC-A was almost eliminated (< 5%ID), whereas that of [^11^C]HC-P remained at over 50%ID. The radioactivity in the C2 compartment representing [^11^C]complex-A or [^11^C]complex-P was approximately 35%ID for both. These results suggest that the reaction between each radioprobe and MAGL reached a saturation dose of approximately 35%ID. More importantly, production of ^11^CO_2_ was detected with [^11^C]HC-A (approximately 35%ID), but not with [^11^C]HC-P (< 1%ID). This result indicates that [^11^C]complex-A was hydrolyzed under the test conditions, resulting in ^11^CO_2_, whereas [^11^C]complex-P was not hydrolyzed.

**Figure 4.**
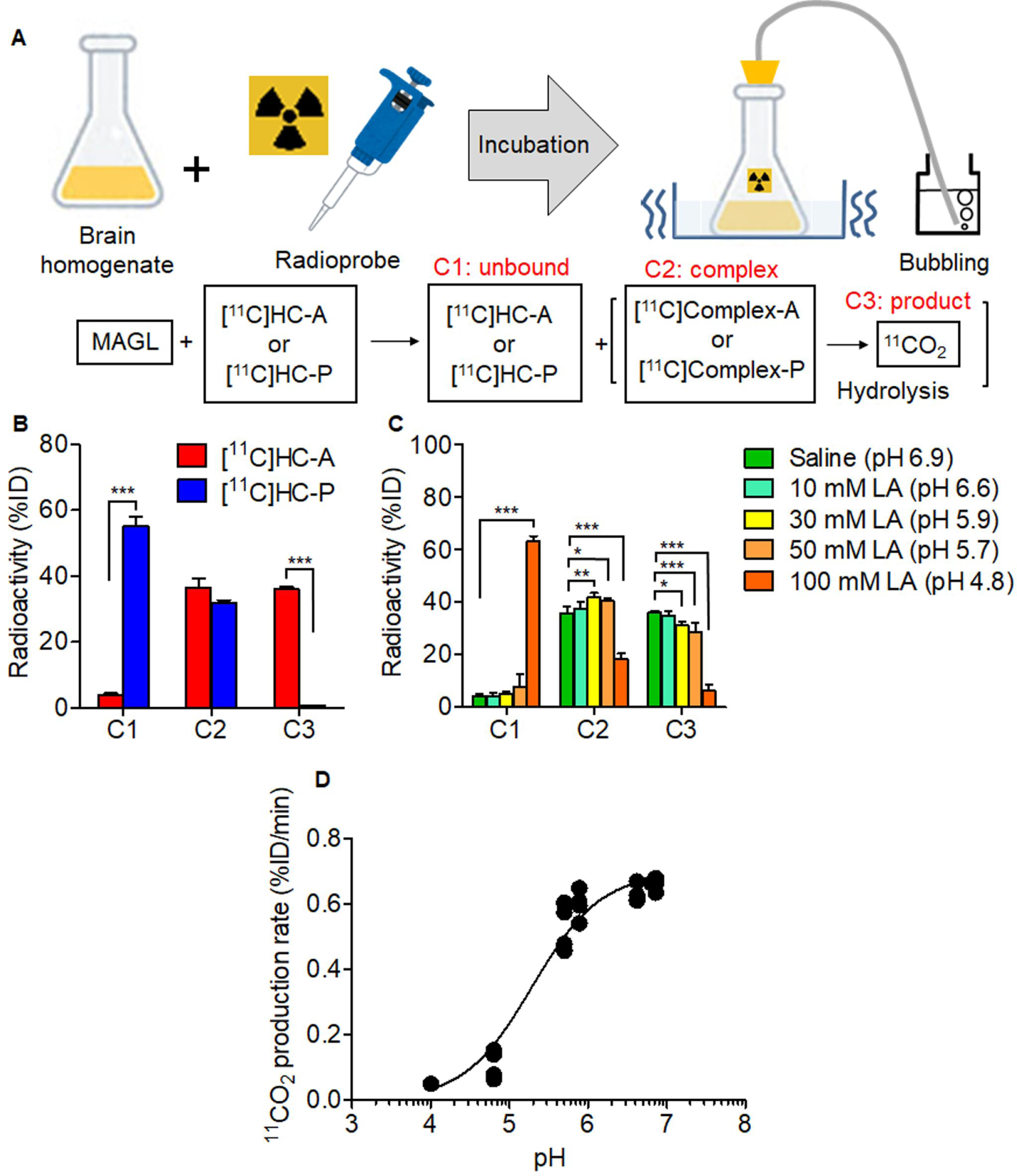
*In vitro* ^11^CO_2_ collection assay using rat brain homogenate. A Schematic illustration and compartments of this assay. B Radioactivity (%ID) derived from [^11^C]HC-A or [^11^C]HC-P in the three divided compartments (C1: unbound; C2: complex; C3: product). Data from three independent experiments for each radioprobe are shown as mean ± SD. \**P* < 0.05, \*\**P* < 0.01, \*\*\**P* < 0.001 (two-way ANOVA with Bonferroni post hoc test). C Radioactivity (%ID) derived from [^11^C]HC-A in the presence of different concentrations (0, 10, 30, 50, and 100 mM) of lactic acid (LA). Data from four independent experiments for each LA concentration are shown as mean ± SD. \**P* < 0.05, \*\**P* < 0.01, \*\*\**P* < 0.001 (two-way ANOVA with Bonferroni post hoc test). D The pH-response curve for ^11^CO_2_ production rate.

We subsequently determined the effect of pH on the rate of ^11^CO_2_ production using an *in vitro* ^11^CO_2_ collection assay with multiple concentrations of lactic acid (LA). Fig 4C shows the radioactivity generated in response to different pH values in each compartment. In the C3 compartment, radioactivity gradually decreased with a lowering of pH. The pH evoking half the maximal ^11^CO_2_ production (pH_50_) was 5.3 (Fig 4D). In contrast, the radioactivity in the C2 compartment was approximately 40%ID with all LA concentrations except for 100 mM (pH 4.8), which resulted in a large decrease in radioactivity, similar to that in the C3 compartment. However, the radioactivity in the C1 compartment after treatment with 100 mM LA increased to > 60%ID. These results suggest that under severely acidic conditions, the ^11^CO_2_ production rate is slow (< pH 4.8) and the binding affinity of [^11^C]HC-A to MAGL is decreased.

In this study, we demonstrated that the hydrolysis of [^11^C]HC-A resulted in ^11^CO_2_ as the final product. Moreover, the production rate of ^11^CO_2_ decreased at lower pH values, which supported the predictions from the MD simulations. Although highly acidic conditions (< pH 4.8) seemed to deactivate MAGL by denaturing it, the binding affinity of [^11^C]HC-A to MAGL and the^11^CO_2_ production were maintained down to pH 5.7. Because the optimum pH for MAGL activity in the brain is in the range of 8–9 (Evagorou *et al*, 2010), this approach can be used to detect activity across a pH range of 5.7–9.

### PET imaging in healthy subject

Prior to the *in vivo* visualization of the pHi gradient using the disease model, we compared the brain kinetics of [^11^C]HC-A and [^11^C]HC-P using PET imaging in healthy rats. Figs 5A–D show representative PET images and time-activity curves (TACs) of [^11^C]HC-A and [^11^C]HC-P across brain regions. PET images of both radioprobes showed widespread radioactive signals throughout the brain. For [^11^C]HC-A, the radioactivity showed a gradual clearance after the initial uptake because [^11^C]complex-A was further hydrolyzed *in vivo*. However, [^11^C]complex-P was not hydrolyzed *in vivo* and the radioactivity remained at the standardized uptake value (SUV) of 1.0–1.4 without clearance at 15 min post-injection. The highest radioactive uptake for [^11^C]HC-A was 1.6–1.9 SUV in each investigated region, which was slightly higher than that of our previously developed [^11^C]compound **6** (1.4–1.7 SUV). This was possibly due to the two-fold higher affinity of [^11^C]HC-A (IC_50_ = 0.2 nM) to MAGL compared with that of [^11^C]compound **6** (IC_50_ = 0.4 nM) (Mori *et al*, 2019).

**Figure 5.**
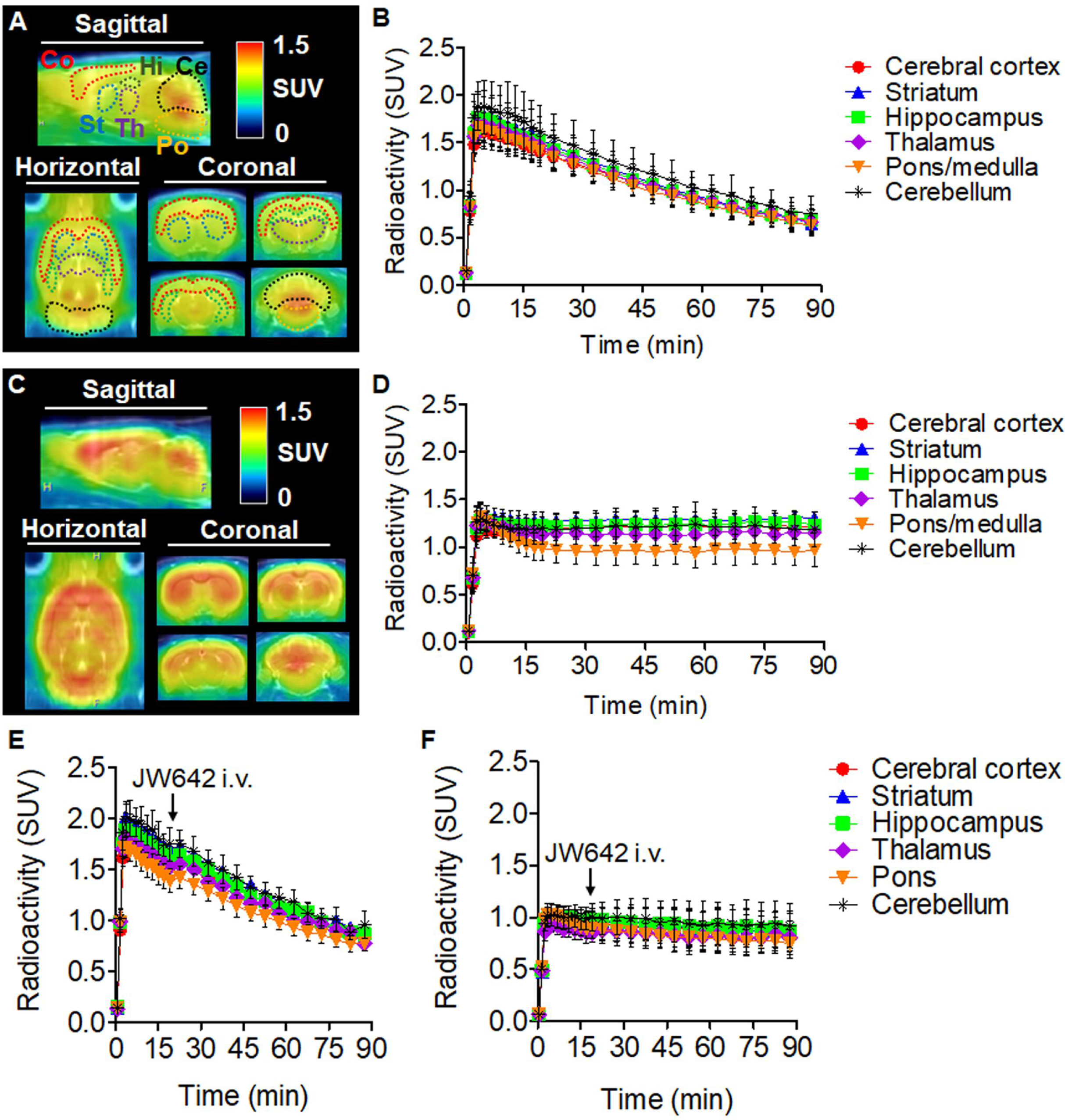
PET imaging in the brain of healthy rat. A–D Averaged PET images and time-activity curves across brain regions of healthy rats. (A) Representative PET/MRI images of [^11^C]HC-A. (B) Time-activity curves (n = 4) of [^11^C]HC-A in the cerebral cortex, striatum, hippocampus, thalamus, pons/medulla, and cerebellum. (C) Representative PET/MRI images of [^11^C]HC-P. (D) Time-activity curves (n = 4) of [^11^C]HC-P in different brain regions. Radioactivity is expressed as SUV. Abbreviations: Co, cerebral cortex; St, striatum; Hi, hippocampus; Th, thalamus; Po, pons/medulla; Ce, cerebellum; SUV, standardized uptake value. E, F Chase studies of [^11^C]HC-A and [^11^C]HC-P using an irreversible-type inhibitor for MAGL (JW642). (E) Time-activity curves (n = 3) of [^11^C]HC-A in the brain of rat treated with JW642 (1 mg/kg, i.v.) 20 min after the scan started. (F) Time-activity curves (n = 3) of [^11^C]HC-P in the brain of rat administered with JW642 (1 mg/kg, i.v.) 20 min after the scan started. Regions of interest were drawn in the cerebral cortex, striatum, hippocampus, thalamus, pons, and cerebellum.

Fig 5E shows chase studies of [^11^C]HC-A and [^11^C]HC-P for determination of the irreversible binding of these radioprobes for MAGL. In this study, the commercial inhibitor JW642 was used as a potent irreversible inhibitor for MAGL because of its structural similarity. Pre-treatment with 1 mg/kg JW642 (i.v.) significantly decreased brain uptake of both radioprobes, the same as self-blocking (Fig EV1). However, there were no changes in the radioactive uptake of either radioprobe when JW642 was post-administered, which strongly suggests that [^11^C]HC-A and [^11^C]HC-P are irreversibly bound to MAGL in brain tissue.

### Theory for quantification of hydrolysis rate

Here, we propose a compartment model (Figs 6A and B) to understand the dynamics of [^11^C]HC-A in the brain, based on the results of the PET studies using healthy rats and the predicted transformation of [^11^C]azetidine carbamate shown in Fig 1A. When [^11^C]HC-A is injected, it enters the bloodstream, reaches the brain capillaries (C_P_), passively crosses (K_1_) the blood–brain barrier (BBB), and enters brain tissue (C_1_). Subsequently, [^11^C]HC-A irreversibly binds (k_3_) to intracellular MAGL to form [^11^C]complex-A (C_2_ compartment), which gradually hydrolyses (K_H_), producing ^11^CO_2_ (C_3_ compartment), and the radioactivity from^11^CO_2_ is rapidly cleared (k_5_) from the brain tissue. Theoretically, when the input function of the radioactivity is close to a negligible value (e.g., 15 min after the injection), the clearance rate of radioactivity in TAC would depend on K_H_ and the efflux rate (k_5_) of ^11^CO_2_. The efflux rate of ^11^CO_2_ has been reported to be 1.74 min^−1^ in the brains of healthy humans (Brooks *et al*, 1984), which is very rapid. Therefore, in our PET experiment, we considered the clearance rate of the radioactivity on PET with [^11^C]HC-A to be the K_H_ value. Here, the clearance rate of the radioactivity is simply estimated by a mono-exponential fitting with a nonlinear least-squares regression (Okamura *et al*, 2009). In this case, the radioactivity clearance rate (≈ K_H_) was 0.01 min^−1^ for all brain regions of a healthy rat.

**Figure 6.**
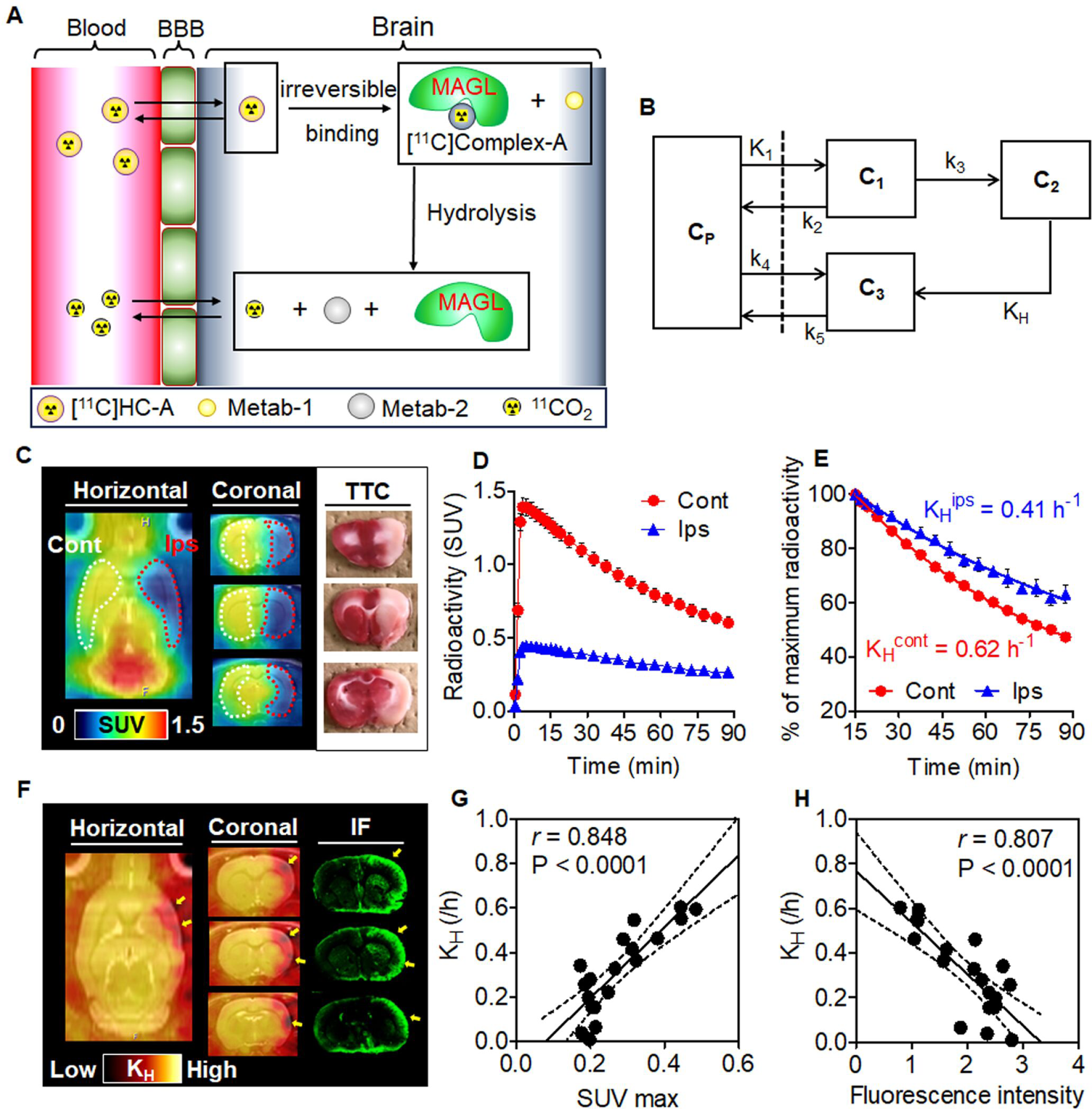
PET imaging in the brain of middle cerebral artery occlusion (MCAO) rat. A, B Model for quantification of the hydrolysis rate (K_H_) of MAGL on PET with [^11^C]HC-A. (A) Diagrammatic reasoning of the clearance of radioactivity starting from [^11^C]HC-A. (B) Schematic of a compartment model for estimation of the kinetic parameters of the radioactivity. C_P_: free radioactive compounds containing [^11^C]HC-A and ^11^CO_2_ in the plasma; C_1_: free and nonspecific binding of [^11^C]HC-A in the brain tissue; C_2_: state under [^11^C]complex-A; C_3_: ^11^CO_2_ as the final product of hydrolysis in the brain tissue; K_1_: influx rate of [^11^C]HC-A; k_2_ efflux rate of [^11^C]HC-A; k_3_: association rate of [^11^C]HC-A to MAGL; K_H_: hydrolysis rate of MAGL; k_4_: reuptake rate of ^11^CO_2_; k_5_: efflux rate of ^11^CO_2_ from the brain side. C–E PET imaging with [^11^C]HC-A in the brains of rats subjected to MCAO for 3–4 hours. (C) Representative PET images and TTC staining. PET images were generated by summing for 0–90 min after injection and scaling with the SUV. The brain slices were prepared post PET scan and stained using TTC. (D) Time–activity curves of [^11^C]HC-A in contralateral (Cont) and ipsilateral (Ips) sides. (E) Clearance rate of radioactivity in contralateral (Cont) and ipsilateral (Ips) sides. Radioactivity is expressed as the percentage of maximum radioactivity (SUV value of 15 min). K_H_ was generated using mono-exponential fitting. F–H *In vivo* mapping of K_H_ levels in MCAO rat. (F) PET images scaled with relative K_H_ value and immunofluorescent (IF) images reflecting hypoxic regions. Yellow arrows indicate areas of intense reduction of K_H_ value and strong fluorescent signals. (G) Correlation plots between K_H_ values and maximum SUV values (SUV max) reflecting cerebral blood flow. (H) Correlation plots between K_H_ values and fluorescence intensity.

### *In vivo* visualization of pHi gradient using PET imaging in brain of experimental stroke model rat

To evaluate the feasibility of *in vivo* visualization of the pHi gradient based on the K_H_ value, we conducted PET experiments with [^11^C]HC-A using middle cerebral artery occlusion (MCAO) rats as a disease model for acute acidic injury resulting from hypoxia. Several reports using MCAO rats showed that lactate concentrations continued to increase in the insult area until 3–4 h after occlusion (Hillered *et al*, 1989; Park *et al*, 2019). Another report using MCAO piglets showed that lactate levels in the insult area peaked within 2–6 h post-occlusion, and then gradually decreased thereafter (Zheng *et al*, 2017). We prepared MCAO rats with different occlusion periods (1, 3–4, or 6 h) and performed PET assessments.

Figs 6C and D show representative averaged PET images of [^11^C]HC-A and TACs on the ipsilateral and contralateral sides of the brains of MCAO rats subjected to 3–4-h occlusion. Radioactive signals disappeared from the ipsilateral side of the forebrain, including the cortex, striatum, and amygdala, which appeared as light areas after 2,3,5-triphenyl tetrazolium chloride (TTC) staining. Additionally, the initial radioactive uptake in the ipsilateral area was approximately 30% of that in the contralateral side. This indicates that the cerebral blood flow (CBF) on the ipsilateral side dropped by 70%. Fig 5E shows a plot of the radioactive clearance in the ipsilateral and contralateral sides of the brains of MCAO rats, including mono-exponential regression analyses. The K_H_ values of [^11^C]complex-A in the ipsilateral and contralateral sides were 0.41 and 0.62 h^−1^, respectively. In MCAO rats with 3–4 h of occlusion, the K_H_ value in the ipsilateral area was 34% lower than that in the contralateral area. Furthermore, the K_H_ values of rats with 1 and 6 h of occlusion were 0.53 and 0.49 h^−1^, respectively, in the ipsilateral sides, and 0.66 and 0.63 h^−1^ in the contralateral sides. The reduced percentage of K_H_ on the ipsilateral side was 25% for 1-h occlusion and 17% for 6-h occlusion (Table 1). These results correspond with previously reported values for the time-dependent changes in intracellular lactate levels in the MCAO model (Hillered *et al*, 1989; Jokivarsi *et al*, 2010; Park *et al*, 2019).

**Table 1.**
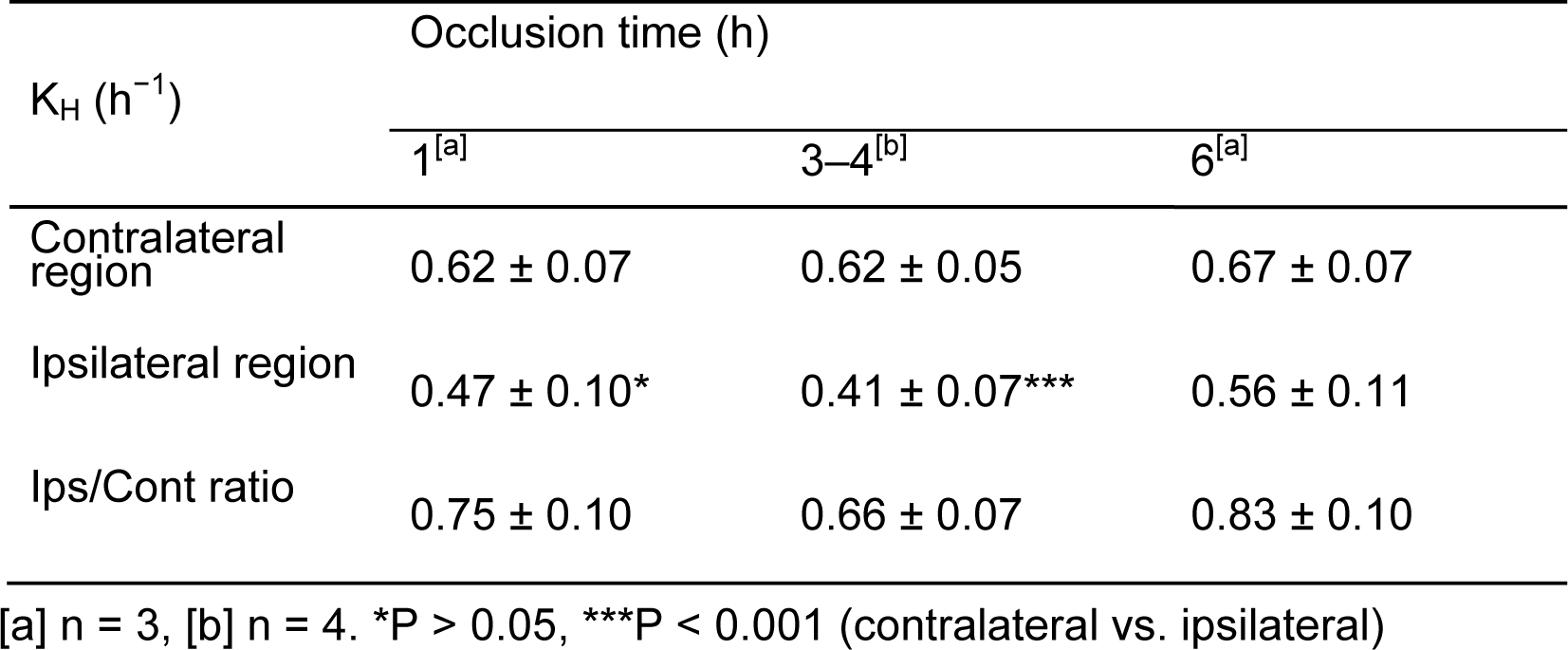
Hydrolysis rate (K_H_) of MAGL in MCAO rats.

We further investigated the effects of different K_H_ values during hypoxia, leading to anaerobic glycolysis and intracellular acidosis. Using the K_H_ value, we performed *in vivo* mapping of the pHi in the brains of MCAO rats with 3–4-h occlusion. Moreover, we compared the K_H_ values in brain regions with immunofluorescent signals from hypoxic areas. As shown in Fig 6F, a low K_H_ value was detected in the upper lip region of the primary somatosensory cortex, secondary somatosensory cortex, insular cortex, and piriform cortex, which correspond to regions with severe reduction in CBF by MCAO (Tsuchidate *et al*, 1997). In these regions, maximum SUV value was also low and exhibited positive correlation to K_H_ value (Fig 6G). Moreover, immunofluorescent signals on slides corresponding to regions of interest in the PET images showed a strong negative correlation (*r* > 0.8, P < 0.0001) with K_H_ values (Fig 6H). Thus, we successfully demonstrated *in vivo* mapping of K_H_ values in the brain, using PET with [^11^C]HC-A to image change in pHi. The MCAO model has also been used for amide proton transfer (APT)-magnetic resonance imaging (MRI), a known pH-weighted imaging technique (Jokivarsi *et al*, 2007; Jokivarsi *et al*, 2010; Zheng *et al*, 2017; Park *et al*, 2019) that is widely used in clinical studies of hypoxia in cases of stroke and brain tumor (Zhou *et al*, 2019). However, a significant correlation between APT signals and intracellular lactate levels has not been found (Zheng & Wang, 2017; Park *et al*, 2019). This might be because of technical issues with MRI, making it suitable for morphological imaging but not for functional imaging. Therefore, our proposed technique, which uses PET imaging, may overcome the low sensitivity to pHi of the APT-MRI technique.

In addition to intracellular enzyme activity (as in the case of MAGL), cytoskeletal component integration, cellular growth, and differentiation rates are all highly affected by pHi. The common characteristics of different CNS diseases such as ischemic stroke, traumatic brain injury, epilepsy, Parkinson’s disease, and Alzheimer’s disease are partly induced by decreases in associated enzymes, which show altered activity under acidic conditions (Fang *et al*, 2010). Furthermore, it is not only ischemia and tumors that cause intracellular acidification, but also the aging process (Bădescu *et al*, 2016). Thus, *in vivo* monitoring of pHi using PET with [^11^C]HC-A could be a promising tool for developing new therapeutic approaches against multiple CNS disorders and for understanding the pathological mechanism of the aging process.

## Materials and Methods

### General

All chemical reagents and organic solvents were purchased from Sigma-Aldrich (St. Louis, MO, USA), FUJIFILM Wako Pure Chemical Industries (Osaka, Japan), or Nacalai Tesque (Kyoto, Japan), and were used without further purification. ^11^C was produced using a cyclotron (CYPRIS HM-18; Sumitomo Heavy Industries, Tokyo, Japan).

^1^H NMR and ^13^C NMR spectra were recorded using a JNM-AL-300 spectrometer (JEOL, Tokyo, Japan) with tetramethylsilane as an internal standard. All chemical shifts (δ) are reported in parts per million (ppm) downfield from the standard. High-resolution mass spectrometry (HRMS) was performed on a JEOL NMS-SX102 spectrometer (JEOL). Column chromatography was performed using a Wakogel C-200 (FUJIFILM Wako Pure Chemical Industries). HPLC was performed on a JASCO HPLC system (JASCO, Tokyo, Japan). Effluent radioactivity was monitored using a NaI (Tl) scintillation detector system. Unless otherwise stated, radioactivity was measured using an IGC-3R Curiemeter (Hitachi Aloka Medical, Tokyo, Japan).

### Animals

Male Sprague-Dawley (SD) rats (n = 24, 7–10 weeks old) weighing 280– 350g were purchased from Japan SLC (Shizuoka, Japan), kept in a temperature- and humidity-controlled environment with a 12-h light-dark cycle, and allowed to freely consume a standard diet (MB-1/Funabashi Farm, Chiba, Japan) and water. All animal experiments were performed according to the recommendations specified by the Committee for the Care and Use of Laboratory Animals of the National Institutes for Quantum and Radiological Science and Technology and Animal Research: Reporting of In Vivo Experiments (ARRIVE) guidelines and were approved by the Institutional Committee of the National Institutes for Quantum Science and Technology (approval number: 16-1006).

### Molecular dynamics (MD) simulations

A three-dimensional (3D) MAGL structure (PDB code: 6AX1), including the ligand-binding domain, was used for modeling and MD simulations. The initial structures of the carbamate inhibitor-MAGL complexes (complex-A and complex-P) were constructed based on model ligands (azetidine carbamate: HC-A; piperidine carbamate: HC-P) in the homology module of the molecular modeling software MOE (Chemical Computing Group, Montreal, Canada). MD simulations for complex-A were conducted under neutral (pH 7) and acidic (pH 6) conditions. Under neutral conditions, Asp and Glu residues and the C-terminus of MAGL were kept deprotonated (COO^−^, Lys, and Arg residues), the N-terminus of the protein were kept in protonation, and the His residues of MAGL were kept in deprotonation. Under acidic conditions, the His residues of MAGL were protonated.

Azetidine-or piperidine-carbamate-MAGL complexes (complex-A and complex-P) were solvated in a truncated octahedral TIP3P water box with a thickness of 8 Å around the complex. Then, protein.ff14SB, water.tip3p, and general amber force field were used as force fields. The systems were neutralized by adding Na^+^ ions as the counter ions. The energy minimized (MM) calculations and MD simulations were performed using AMBER18/PMEMD with periodic boundary conditions of constant temperature (300 K) and pressure (1 atm) and a non-bonded interaction cutoff distance of 8 Å. The particle mesh Ewald method was employed to calculate the electrostatic interactions. After tethering the heavy atoms of the protein, the MM calculations of the system were performed in 1000 steps using the steepest descent method and the conjugated gradient method. MD simulations were performed only for water molecules in the system in a 50 ps time frame while increasing the temperature from 0 to 300 K. Subsequently, the Cα atoms of the protein were tethered, then MM calculations of the side chain of the protein were performed in 1000 steps using the steepest descent method and in 10 000 steps using the conjugated gradient method. MD calculations for the Cα atoms of the protein were performed under the same conditions described above. MM calculations for the system were performed in 1000 steps using the steepest descent method and in 10 000 steps using the conjugated gradient method. Finally, the MM calculations and MD simulations were performed for the entire system. Atoms were heated from 0 to 300 K at 50 ps intervals and the whole system reached the equilibrium state for MD simulation (5 ns) of the complex structures under constant temperature (300 K) and pressure (1 atm). Structural changes were evaluated using the root mean square deviation of the trajectory of the intermediate structures in the MD simulations for each complex.

### Chemistry

#### Synthesis of 1,1,1,3,3,3-hexafluoropropan-2-yl 3-(1-phenyl-1H-pyrazol-3-yl)azetidine-1-carbamate (HC-A)

HC-A was synthesized according to a reported method (Jones *et al*, 2016; Butler *et al*, 2017), with some modifications as shown in Fig EV2A. Briefly, tert-butyl 3-(1-phenyl-1*H*-pyrazol-3-yl)azetidine-1-carboxylate was synthesized by reacting tert-butyl 3-(1*H*-pyrazol-3-yl)azetidine-1-carboxylate (314 mg, 1.4 mmol) with a mixture of phenylboronic acid (343 mg, 2.8 mmol), Cu(OAc)_2_ (383 mg, 2.1 mmol), pyridine (227 μL, 2.8 mmol), and CH_2_Cl_2_ (3 mL) under N_2_. 3-(Azetidin-3-yl)-1-phenyl-1*H*-pyrazole (precursor A) was synthesized via a deprotection reaction of tert-butyl 3-(1-phenyl-1*H*-pyrazol-3-yl)azetidine-1-carboxylate with HCl (4 mol/L 1,4-dioxane, 5.8 mL). 1,1,1,3,3,3-Hexafluoropropan-2-ol (22 μL, 0.2 mmol) in anhydrous CH_2_Cl_2_ (1 mL) was mixed with DIPEA (45 μL, 0.2 mmol) and DMAP (2.8 mg, 0.02 mmol) and then reacted with triphosgene (21 mg, 0.07 mmol) in CH_2_Cl_2_ (1 mL) at room temperature (r.t.) under N_2_. The reaction mixture was then stirred overnight. 4-(1-Phenyl-1*H*-pyrazol-3-yl)azetidine (50 mg, 0.2 mmol) and DIPEA (120 μL, 0.6 mmol) in CH_2_Cl_2_ (1 mL) were added to the mixture and stirred at r.t. for 5 h. The solution was evaporated, extracted with AcOEt and water, and dried using Na_2_SO_4_. The filtrate was concentrated under vacuum and the obtained crude product was purified using silica gel column chromatography (hexane/ethyl acetate= 6/1) to give HC-A (27 mg, 33%) as a colorless oil. ^1^H NMR (300 MHz, CDCl_3_) 7.90 (d, J = 2.7 Hz, 1H), 7.66 (d, J = 8.4 Hz, 2H), 7.46 (t, J = 7.2, 8.4 Hz, 2H), 7.29 (t, J = 7.8, 7.2 Hz, 1H), 6.41 (d, J = 2.4 Hz, 1H), 5.75–5.63 (m, 2H), 4.55–4.46 (m, 2H), 4.37–4.29 (m, 2H), 4.09–4.01 (m, 1H). HRMS (ESI): calculated m/z for C_16_H_14_O_2_N_3_F_6_ [M+H]^+^: 394.0990; found 394.0999.

The analytical data were identical to the reported data (Jones *et al*, 2016; Butler *et al*, 2017).

#### Synthesis of tert-butyl 4-(1-phenyl-1H-pyrazol-3-yl)piperidine-1-carboxylate

*(E)*-Tert-butyl 4-(3-(dimethylamino)acryloyl)piperidine-1-carboxylate and tert-butyl 4-(1*H*-pyrazol-3-yl)piperidine-1-carboxylate were synthesized according to a reported method (Butler *et al*, 2017) as shown in Fig EV2B. Briefly, tert-butyl 4-acetylpiperidine-1-carboxylate (227 mg, 1 mmol) was reacted with DMF-DMA (1.0 mL, 8 mmol) under reflux for 18 h. The reaction mixture was diluted with toluene (1.0 mL) and evaporated to yield *(E)*-Tert-butyl 4-(3-(dimethylamino)acryloyl)piperidine-1-carboxylate as a brown oil. Subsequently, crude product of *(E)*-Tert-butyl 4-(3-(dimethylamino)acryloyl)piperidine-1-carboxylate was dissolved in EtOH (6 mL), which was reacted with hydrazine hydrate (75 μL, 1.5 mmol) under reflux for 14.5 h. Removal of EtOH yielded tert-butyl 4-(1*H*-pyrazol-3-yl)piperidine-1-carboxylate as a brown oil without further purification. Crude product of tert-butyl 4-(1*H*-pyrazol-3-yl)piperidine-1-carboxylate (251 mg, 1 mmol) was reacted with the mixture containing phenylboronic acid (244 mg, 2 mmol), Cu(OAc)_2_ (272.2 mg, 1.5 mmol), pyridine (161 µL, 2 mmol), and CH_2_Cl_2_ (3 mL) under N_2_ at r.t. for 12 h. The reaction mixture was filtered through celite and the filtrate was concentrated by evaporation. The crude product was purified as a light-yellow solid using silica gel column chromatography (hexane/ethyl acetate= 6/1) to give tert-butyl 4-(1-phenyl-1*H*-pyrazol-3-yl)piperidine-1-carboxylate (272 mg, 83%).^1^H NMR (300 MHz, CDCl_3_) 7.83 (d, J = 2.4 Hz, 1H), 7.65 (d, J = 8.1 Hz, 2H), 7.43 (t, J = 8.1, 8.1 Hz, 2H), 7.27–7.22 (m, 1H), 6.27 (d, J = 2.7 Hz, 1H), 4.26–4.08 (m, 2H), 2.95–2.84 (m, 3H), 2.02–1.97 (m, 2H), 1.75–1.61 (m, 2H), 1.48 (s, 9H).

#### Synthesis of 4-(1-phenyl-1H-pyrazol-3-yl)piperidine (precursor B)

Tert-butyl 4-(1-phenyl-1*H*-pyrazol-3-yl)piperidine-1-carboxylate (271 mg, 0.83 mmol) was treated with HCl (4 mol/L in 1,4-dioxane, 4 mL) and the reaction mixture was stirred at r.t. for 1 h. Removal of the solvents yielded 4-(1-phenyl-1*H*-pyrazol-3-yl)piperidine (precursor B, 140 mg, 75%) as a colorless solid. ^1^H NMR (300 MHz, CDCl_3_) 7.86 (d, J = 2.4 Hz, 1H), 7.65 (d, J = 8.1 Hz, 2H), 7.45 (t, J = 7.8, 7.8 Hz, 2H), 7.32–7.29 (m, 1H), 6.34 (d, J = 2.7 Hz, 1H), 3.76–3.70 (m, 2H), 3.63–3.55 (m, 3H), 3.25–3.10 (m, 2H), 2.35–2.07 (m, 2H). HRMS (ESI): calculated m/z for C_14_H_18_N_3_ [M+H]^+^: 228.1501; found 228.1463.

#### Synthesis of 1,1,1,3,3,3-hexafluoropropan-2-yl-4-(1-Phenyl-1H-pyrazol-3-yl)piperidine-1-carboxylate (HC-P)

A solution of 1,1,1,3,3,3-hexafluoropropan-2-ol (26 μL, 0.2 mmol) in anhydrous CH_2_Cl_2_ (1.5 mL) was reacted with di-2-pyridylcarbonate (52 mg, 0.2 mmol) and DMAP (3.0 mg, 0.02 mmol) at r.t. under N_2_ overnight with stirring. Precursor B (50 mg, 0.22 mmol), DIPEA (115 μL, 0.66 mmol), and CH_2_Cl_2_ (1.0 mL) were added to the mixture and then stirred at r.t. overnight. After the solution had evaporated, the crude product was purified by silica gel column chromatography (hexane/ethyl acetate= 8/1) to give HC-P (21.0 mg, 23%) as a colorless oil. ^1^H NMR (300 MHz, CDCl_3_) 7.84 (d, J = 2.4 Hz, 1H), 7.65 (d, J = 8.1 Hz, 2H), 7.44 (t, J = 7.5, 8.0 Hz, 2H), 7.29–7.24 (m, 1H), 6.27 (d, J = 2.7 Hz, 1H), 5.84–5.72 (m, 1H), 4.26–4.18 (m, 2H), 3.18–3.09 (m, 2H), 3.05–2.97 (m, 1H), 2.11–2.04 (m, 2H), 1.85–1.7 (m, 2H). HRMS (ESI): calculated m/z for C_18_H_18_F_6_N_3_O_2_ [M+H]^+^: 422.1303; found 422.1327.

### Radiochemistry

^11^CO_2_ was produced via ^14^N(p, α)^11^C nuclear reactions using a cyclotron (Cypris HM18, Sumitomo Heavy Industries) and transferred into a pre-heated mechanizer packed with a nickel catalyst at 400°C to produce ^11^CH_4_, which was subsequently reacted with chlorine gas at 560°C to generate ^11^CCl_4_. ^11^COCl_2_ was produced via the reaction of ^11^CCl_4_ with iodine oxide and sulfuric acid and trapped in a solution of 1,1,1,3,3,3-hexafluoropropan-2-ol (3.1 μL) and 1,2,2,6,6-pentamethylpiperidine (PMP; 5.4 μL) in THF (200 µL) at 0°C. The solution was then heated to 30°C for 3 min. A solution of precursor A or B (1.00 mg) with PMP (2.3 µL) in THF (200 µL) was added to the mixture and heated at 30°C for 3 min and the solvent was then removed at 80°C before cooling to room temperature. After dilution with 1 mL of separation solvent, the mixture was purified by HPLC (CAPCELL Pak C18 column; 10 mm i.d. × 250 mm, 5 µm) using a mobile phase of CH_3_CN/H_2_O/Et_3_N (70/30/0.1, v/v/v, for [^11^C]HC-A) or CH_3_CN/H_2_O/TFA (75/25/0.1, v/v/v, for [^11^C]HC-P) at a flowrate of 5.0 mL/min. The respective retention times were 11.0 min for [^11^C]HC-A and 11.7 min for [^11^C]HC-P. The HPLC fractions of [^11^C]HC-A and [^11^C]HC-P were collected in a flask containing 25% ascorbic acid in sterile water (100 µL), and Tween 80 (75 µL) in ethanol (0.3 mL) was added before synthesis and evaporation to dryness. The final product was collected in a sterile flask and reformulated in sterile normal saline (2 mL). The resulting compound was analyzed using HPLC with a UV detector of 254-nm wavelength and a radioactivity detector. The radiochemical and chemical purities were measured by analytical HPLC (CAPCELL Pak C18 column; 4.6 mm i.d. × 250 mm, 5 µm) as a mobile phase of CH_3_CN/H_2_O/Et_3_N (70/30/0.1, v/v/v, for [^11^C]HC-A) or CH_3_CN/H_2_O/TFA (75/25/0.1, v/v/v, for [^11^C]HC-P) at a flow rate of 1.0 mL/min. The identification of [^11^C]HC-A and [^11^C]HC-P was confirmed by co-injection with HC-A or HC-P, respectively, and the retention times in radio-HPLC analyses were 10.3 min for [^11^C]HC-A and 10.8 min for [^11^C]HC-P. The calculated lipophilicities (cLogP) of each radioligand were estimated by ChemBioDraw Ultra (version 12.0; Perkin Elmer, Waltham, MA, USA).

### *In vitro* ^11^CO_2_ collection assay

Six rats were anesthetized with isoflurane (1–5% in air) and exsanguinated. The brain was quickly removed and homogenized with three equivalents of saline using a homogenizer (Silent Crasher S; Heidolph, Schwabach, Germany). The homogenate (6 mL) was added to a conical flask containing saline (2 mL) and lactic acid (1 mL) at various concentrations (0, 10, 30, 50, and 100 mM). The flask was closed with rubber plug inserting glass tubes to allow bubbling. [^11^C]HC-A (1 mL, 10–18 MBq) or [^11^C]HC-P (1 mL, 10–11 MBq) was added to the flask and the flask was incubated in a 37°C water bath with shaking for 60 min. During the incubation, a tube was inserted into the bottle containing 10 mL of 2-aminoethanol (50% in distilled water). The air for bubbling was then automatically browsed using an air pump connected to another tube. The bottle was then changed for a new one after 5 min. Before stopping the enzymatic reaction, the pH of the brain homogenates was measured using a pH meter (Elite pH Spear Pocket Testers; Thermo Fisher Scientific, Waltham, MA, USA). Subsequently, pre-chilled CH_3_CN (10 mL) was quickly added to the flask to stop the enzymatic reaction and the mixture was separated into several micro-tubes. The micro-tubes were centrifuged at 10 000 *g* and the supernatants were transferred into new micro-tubes. The radioactivity levels of the bottle for collecting ^11^CO_2_, the pellet, and the supernatant were measured using an autogamma scintillation counter (2480 WIZARD^2^; PerkinElmer). Radioactivity is expressed as a percentage of the incubation dose (%ID).

### Permanent middle cerebral artery occlusion (pMCAO) surgery

The pMCAO model was established in 10 rats. The rats were anesthetized with isoflurane (1.5% in air) and their body temperature was maintained at approximately 37°C using a heating pad (BWT-100; Bio Research Center Co., Ltd., Aichi, Japan). The pMCAO surgery was performed using the Koizumi method (Koizumi *et al*, 1986). Briefly, using a surgical microscope (CLS 150MR; Leica Microsystems, Wetzlar, Germany), a silicon-coated monofilament (Doccol Corporation, Sharon, MA, USA) was introduced into the internal carotid artery via a cut in the external carotid artery until the monofilament occluded the MCA base. After 1, 3–4, or 6 h of occlusion, the rats were used for the PET imaging study.

### Small-animal PET imaging

Before PET assessment, the rats were anesthetized with isoflurane (5% in air) and a 24-gauge intravenous catheter (Terumo Medical Products, Tokyo, Japan) was inserted into their tail vein. For the blocking study, unlabeled HC-A and HC-P and JW642 (MedChemExpress, Monmouth Junction, NJ, USA) in different doses (1 or 3 mg/kg) were administered via the tail vein catheter 30 min before the injection of the radioprobe. Subsequently, the rats were secured in a custom-designed chamber and placed in a small-animal PET scanner (Inveon; Siemens Medical Solutions, Knoxville, TN, USA). Their body temperatures were maintained using a 40°C water circulation system (T/Pump TP401; Gaymar Industries, Orchard Park, NY, USA). A bolus of [^11^C]HC-A (1 mL, 52–57 MBq, 0.3–0.9 nmol) or [^11^C]HC-P (1 mL, 41–60 MBq, 1.0–1.2 nmol) was injected at a flow rate of 0.5 mL/min via the tail vein catheter. For the chase study, JW642 at 1 mg/kg was administered via the tail vein catheter 20 min after a bolus injection of radioprobe. Dynamic emission scanning in 3D list mode was performed for 90 min (1 min × 4 frames, 2 min × 8 frames, 5 min × 14 frames). The acquired dynamic PET images were reconstructed by filtered back-projection using a Hanning filter with a Nyquist cutoff of 0.5 cycles/pixel. The time-activity curves (TACs) of [^11^C]HC-A and [^11^C]HC-P were acquired for volumes of interest (VOIs) in the cerebral cortex, striatum (caudate/putamen), hippocampus, thalamus, pons, and cerebellum by referring to a rat brain MRI template using PMOD software (version 3.4; PMOD technology, Zurich, Switzerland). For the MCAO model, VOIs were drawn on both the ipsilateral and contralateral sides of the brain. The radioactivity was decay-corrected to the injection time and is expressed as SUV. The rate of hydrolysis (K_H_) was estimated by a one-exponential fitting on the TACs from 15 to 90 min, as follows:

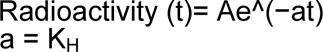

*In vivo* mapping scaled with the relative K_H_ value was generated by dividing 0 to 15-min average images by 15 to 90-min average images using PMOD software.

### TTC staining

After PET scanning, the rats were euthanized by cervical dislocation. The brain was quickly removed, placed on a brain slicer (EMJapan Co., Ltd., Tokyo, Japan), and cut into 2-mm coronal sections. The sections were incubated in 2% TTC (dissolved in phosphate-buffered saline) for 20 min at r.t.

### Immunofluorescent staining for hypoxia

Tissue hypoxia levels were assessed using Hypoxyprobe™ (Hypoxyprobe Inc, Burlington, MA, USA) immunofluorescence staining (Wang *et al*, 2021). Briefly, after occlusion for 4 h in MCAO rats, Hypoxyprobe™ (60 mg/kg body weight) was injected intraperitoneally. The rats were sacrificed after a PET scan (90 min) and their brains were quickly removed and frozen on powdered dry ice. Coronal brain sections were cut at −20°C using a cryostat (NX50; Thermo Fisher Scientific, Waltham, MA, USA) and mounted on MAS-coated glass slides (Matsunami-glass, Osaka, Japan). Cryo-sections were prepared with a 20-μm thickness and a 100-μm distance between two adjacent sections. For the assay, brain sections were fixed in 4% paraformaldehyde/PBS for 15 min at r.t. and then blocked with blocking serum for 20 min at r.t. The sections were incubated with anti-hypoxyprobe mouse IgG as the primary antibody for 30 min at r.t. After thorough washing, the sections were incubated with a biotinylated secondary antibody (1:100) for 30 min at r.t., followed by fluorescent staining using a TSA fluorescence system (PerkinElmer). The slides were then mounted with medium (Vector Laboratories, Burlingame, CA, USA). The fluorescent signals and areas with fluorescent staining in regions of interest (ROIs) shown in Fig EV3 were quantified using ImageJ software. The fluorescence intensities were compared with K_H_ values obtained from the same individual in the PET study using [^11^C]HC-A.

### Statistical Analysis

Statistical analyses were performed using GraphPad Prism (version 5.0; GraphPad Software, San Diego, CA, USA). Differences between means were assessed as appropriate, using two-way ANOVA followed by Bonferroni post-hoc analysis.

## Data availability

This study contains no data deposited in external repositories. Expanded View for this article is available online.

## Acknowledgements

We thank the staff of the National Institutes for Quantum and Radiological Science and Technology for their support with the following: cyclotron operation, radioisotope production, radiosynthesis, and animal experiments. We thank Edanz (https://jp.edanz.com/ac) for editing a draft of this manuscript. This work was supported by QST President’s Strategic Grant (Exploratory Research to TY), JSPS KAKENHI (Grant No. 20H03635 to MRZ), and the AMED Moonshot Research and Development Program (Grant No. 21zf0127003h001 to MRZ).

## Author contributions

TY, WM, and TO were contributions equally. TY and MRZ were responsible authors for this work. The manuscript was written by TY with contributions of WM, TO, and MRZ. Funding for this work was acquired by TY and MRZ. Chemistry and radiochemistry were primary contributed by WM with supports of YK, NN, and MF. Molecular docking simulation study was conducted by TO. All animal experiments were mainly performed by AH with helping of YZ and TY. Development of kinetic model and data analysis were conducted by TY with supports of HT. All authors have read and agreed to the published version of the manuscript.

## Conflict of interest

The authors declare that they have no conflict of interest.

## The Paper Explained

### Problem

Although intracellular accumulation of acidic sources was harmful for neurons and know to be involved in multiple CNS disorders, there were no PET probes to monitor intracellular pH *in vivo*.

## Results

MD simulations predicted that HC-A, a covalent MACL inhibitor, followed by forming complex with MAGL, was easily hydrolyzed *in vivo*. However, under weakly acidic conditions (pH 6.0), the complex was stable against hydrolysis probably due to a change in the spatial arrangement of the water molecules. Moreover, *in vitro* assays with [^11^C]HC-A using rat brain homogenate also showed a pH-dependent reduction in the hydrolysis rate of [^11^C]complex. Taking advantage of this property, we developed to estimate *in vivo* hydrolysis rate (K_H_) of [^11^C]complex by using monoexponential fitting of time-activity curves of [^11^C]HC-A in the ipsilateral or contralateral brain regions of MCAO rats. Further, it was demonstrated that the K_H_ value on the ipsilateral side decreased with a peak at 3–4 h occlusion, which corresponded with the time-course change in the intracellular concentration of lactate. Thus, we successfully developed a new PET imaging technique scaled with K_H_ corresponding to changes in intracellular pH in the brain using [^11^C]HC-A.

### Impact

Our proposed PET imaging technique for *in vivo* mapping intracellular pH gradient in brain would permits furtherprogressing diagnostic studies and furtherunderstanding of mechanisms in multiple CNS disorders, gliomas, and aging.

## Expanded View Figure legends

**Figure EV1. Blocking studies for PET with [^11^C]HC-A and [^11^C]HC-P.**

A Time-activity curves of [^11^C]HC-A in the whole brain of rat administrated with or without HC-A or JW642 of different doses (1 and 3 mg/kg). The blockade by JW642 made more reduction than that by HC-A. These differences might be caused by overflows from peripheral organs by self-blocking. There were no differences between different doses of both blockers.

B Time-activity curves of [^11^C]HC-P in the whole brain of rat administrated with or without HC-P or JW642 of different doses (1 and 3 mg/kg). Although intensive increments of radioactive uptake in initial phase were seen in subjects treated with JW642, radioactivity in the brain of rat treated with blocking agent significantly subsequently decreased to 0.5 SUV. There were no differences between different doses of both blockers.

**Figure EV2. Chemical syntheses of HC-A and HC-P.**

A Scheme of chemical synthesis of HC-A. HC-A was synthesized by reacting with precursor A produced starting from tert-butyl 3-(1*H*-pyrazol-3-yl)azetidine-1-calboxylate.

B Scheme of chemical synthesis of HC-P. HC-P was synthesized by reacting with precursor B derived from tert-butyl 4-(1-phenyl-1*H*-pyrazol-3-yl)piperidine-1-carboxylate.

**Figure EV3. Regions of interest (Rois) in images of immunofluorescence staining corresponding to PET images of MCAO rat.**

The Rois were manually drawn on the top of striatum (Roi 1), bottom of striatum (Roi 2), primary somatosensory cortex (Roi 3), secondary somatosensory cortex (Roi 4), insular cortex (Roi 5), and piriform cortex (Roi 6) of ipsilateral hemisphere. The Roi on the striatum of contralateral hemisphere is described as the reference.

## Notes

### Competing Interest Statement

The authors have declared no competing interest.

## References

Bădescu GM, Fîlfan M, Ciobanu O, Dumbravă DA, Popa-Wagner A (2016) Age-related hypoxia in CNS pathology. Rom J Morphol Embryol 57: 33–43

Baillieul S, Chacaroun S, Doutreleau S, Detante O, Pépin JL, Verges S (2017) Hypoxic conditioning and the central nervous system: A new therapeutic opportunity for brain and spinal cord injuries? Exp Biol Med (Maywood) 242: 1198–1206

Barzagli F, Mani F, Peruzzini M (2016) A Comparative Study of the CO_2_ Absorption in Some Solvent-Free Alkanolamines and in Aqueous Monoethanolamine (MEA). Environ Sci Technol 50: 7239–7246

Bouzat P, Oddo M (2014) Lactate and the injured brain: friend or foe? Curr Opin Crit Care 20: 133–140

Brooks DJ, Lammertsma AA, Beaney RP, Leenders KL, Buckingham PD, Marshall J, Jones T (1984) Measurement of regional cerebral pH in human subjects using continuous inhalation of 11CO2 and positron emission tomography. J Cereb Blood Flow Metab 4: 458–465

Butler CR, Beck EM, Harris A, Huang Z, McAllister LA, Am Ende CW, Fennell K, Foley TL, Fonseca K, Hawrylik SJ, et al (2017) Azetidine and Piperidine Carbamates as Efficient, Covalent Inhibitors of Monoacylglycerol Lipase. J Med Chem 60: 9860–9873.

Cheng R, Mori W, Ma L, Alhouayek M, Hatori A, Zhang Y, Ogasawara D, Yuan G, Chen Z, Zhang X, et al (2018) In Vitro and in Vivo Evaluation of 11C-Labeled Azetidinecarboxylates for Imaging Monoacylglycerol Lipase by PET Imaging Studies. J Med Chem 61: 2278–2291

Chen Z, Mori M, Deng X, Cheng R, Ogasawara D, Zhang G, Schafroth MA, Dahl K, Fu H, Hatori A, et al (2019) Design, Synthesis, and Evaluation of Reversible and Irreversible Monoacylglycerol Lipase Positron Emission Tomography (PET) Tracers Using a “Tail Switching” Strategy on a Piperazinyl Azetidine Skeleton. J Med Chem 62: 3336–3353

Chesler M (2005) Failure and function of intracellular pH regulation in acute hypoxic-ischemic injury of astrocytes. Glia 50: 398–406

Di Marzo V, Stella N, Zimmer A (2015) Endocannabinoid signalling and the deteriorating brain. Nat Rev Neurosci 16: 30–42

Evagorou A, Anagnostopoulos D, Farmaki E, Siafaka-Kapadai A (2010) Hydrolysis of 2-arachidonoylglycerol in Tetrahymena thermophila. Identification and partial characterization of a Monoacylglycerol Lipase-like enzyme. Eur J Protistol 46: 289–297

Fang B, Wang D, Huang M, Yu G, Li H (2010) Hypothesis on the relationship between the change in intracellular pH and incidence of sporadic Alzheimer’s disease or vascular dementia. Int J Neurosci 120: 591–595

Freeman RS, Barone MC (2005) Targeting hypoxia-inducible factor (HIF) as a therapeutic strategy for CNS disorders. Curr Drug Targets CNS Neurol Disord 4: 85–92

Hillered L, Hallström A, Segersvärd S, Persson L, Ungerstedt U (1989) Dynamics of extracellular metabolites in the striatum after middle cerebral artery occlusion in the rat monitored by intracerebral microdialysis. J Cereb Blood Flow Metab 9: 607–616

Jokivarsi KT, Gröhn HI, Gröhn OH, Kauppinen RA (2007) Proton transfer ratio, lactate, and intracellular pH in acute cerebral ischemia. Magn Reason Med 57: 647–653

Jokivarsi KT, Hiltunen Y, Tuunanen PI, Kauppinen RA, Gröhn OH (2010) Correlating tissue outcome with quantitative multiparametric MRI of acute cerebral ischemia in rats. J Cereb Blood Flow Metab 30: 415–427

Jones A, Kemp M, Stockley M, Gibson K, Whitlock G (2016) 1-CYANO-PYRROLIDINE COMPOUNDS AS USP30 INHIBITORS. WO 2016/156816 A1

Kamimura K, Nakajo M, Yoneyama T, Takumi K, Kumagae Y, Fukukura Y, Yoshiura T (2019) Amide proton transfer imaging of tumors: theory, clinical applications, pitfalls, and future directions. Jpn J Radiol 37: 109–116

Kearfott KJ, Junck L, Rottenberg DA (1983) C-11 dimethyloxazolidinedione (DMO): biodistribution, radiation absorbed dose, and potential for PET measurement of regional brain pH: concise communication. J Nucl Med 24: 805–811

Koizumi J, Yoshida Y, Nakazawa T, Ooneda G (1986) Experimental studies of ischemic brain edema, I: a new experimental model of cerebral embolism in rats in which recirculation can be introduced in the ischemic area. Jpn J Stroke 8: 8

Mori W, Hatori A, Zhang Y, Kurihara Y, Yamasaki T, Xie L, Kumata K, Hu K, Fujinaga M, Zhang MR (2019) Radiosynthesis and evaluation of a novel monoacylglycerol lipase radiotracer: 1,1,1,3,3,3-hexafluoropropan-2-yl-3-(1-benzyl-1H-pyrazol-3-yl)azetidine-1-[11C]carboxylate. Bioorg Med Chem 27: 3568-3573

Okamura T, Kikuchi T, Okada M, Toramatsu C, Fukushi K, Takei M, Irie T (2009) Noninvasive and quantitative assessment of the function of multidrug resistance-associated protein 1 in the living brain. J Cereb Blood Flow Metab 29: 504–511

Park JE, Jung SC, Kim HS, Suh JY, Baek JH, Woo CW, Park B, Woo DC (2019) Amide proton transfer-weighted MRI can detect tissue acidosis and monitor recovery in a transient middle cerebral artery occlusion model compared with a permanent occlusion model in rats. Eur Radiol 29: 4096–4104

Tsuchidate R, He QP, Smith ML, Siesjö BK (1997) Regional cerebral blood flow during and after 2 hours of middle cerebral artery occlusion in the rat. J Cereb Blood Flow Metab 17: 1066–1073.

Uria-Avellanal C, Robertson NJ (2014) Na^+^/H^+^ exchangers and intracellular pH in perinatal brain injury. Transl Stroke Res 5: 79–98

Vāvere AL, Biddlecombe GB, Spees WM, Garbow JR, Wijesinghe D, Andreev OA, Engelman DM, Reshetnyak YK, Lewis JS (2009) A novel technology for the imaging of acidic prostate tumors by positron emission tomography. Cancer Res 69: 4510–4516

Viader A, Blankman JL, Zhong P, Liu X, Schlosburg JE, Joslyn CM, Liu QS, Tomarchio AJ, Lichtman AH, Selley DE, et al (2015) Metabolic Interplay between Astrocytes and Neurons Regulates Endocannabinoid Action. Cell Rep 12: 798–808

Wang E, Wu Y, Cheung JS, Igarashi T, Wu L, Zhang X, Sun PZ (2019) Mapping tissue pH in an experimental model of acute stroke - Determination of graded regional tissue pH changes with non-invasive quantitative amide proton transfer MRI. Neuroimage 191: 610–617

Wang L, Mori W, Cheng R, Yui J, Hatori A, Ma L, Zhang Y, Rotstein BH, Fujinaga M, Shimoda Y, et al (2016) Synthesis and Preclinical Evaluation of Sulfonamido-based [11C-Carbonyl]-Carbamates and Ureas for Imaging Monoacylglycerol Lipase. Theranostics 6: 1145–1159

Wang L, Ren C, Li Y, Gao C, Li N, Li H, Wu D, He X, Xia C, Ji X (2021) Remote ischemic conditioning enhances oxygen supply to ischemic brain tissue in a mouse model of stroke: Role of elevated 2,3-biphosphoglycerate in erythrocytes. J Cereb Blood Flow Metab 41: 1277–1290

Zhang X, Lin Y, Gillies RJ (2010) Tumor pH and its measurement. J Nucl Med 51: 1167–1170

Zheng Y, Wang XM (2017) Measurement of Lactate Content and Amide Proton Transfer Values in the Basal Ganglia of a Neonatal Piglet Hypoxic-Ischemic Brain Injury Model Using MRI. AJNR Am J Neuroradiol 38: 827–834

Zhou J, Heo HY, Knutsson L, van Zijl PCM, Jiang S (2019) APT-weighted MRI: Techniques, current neuro applications, and challenging issues. J Magn Reson Imaging 50, 347–364

